# The mechanics underpinning non-deterministic computation in cortical neural networks

**DOI:** 10.1101/2022.12.03.518983

**Authors:** Elizabeth A Stoll

**Affiliations:** Western Institute for Advanced Study, Denver, USA

## Abstract

Cortical neurons allow random electrical noise to contribute to the likelihood of firing a signal. Previous approaches have involved statistically modeling signaling outcomes in neuronal populations, or modeling the dynamical relationship between membrane potential, ion channel activation, and ion conductance in individual neurons. However, these methods do not mechanistically account for the role of random electrical noise in gating the action potential. Here, the membrane potential of a cortical neuron is modeled as the uncertainty in all component pure states, or the amount of information encoded by that computational unit. With this approach, each neuron computes the probability of transitioning from an off-state to an on-state, with the macrostate of each computational unit being a function of all component microstates. Component pure states are integrated into a physical quantity of information, and the derivative of this high-dimensional probability density yields eigenvalues, or an internally-consistent observable system state at a defined point in time. In accordance with the Hellman-Feynman theorem, the resolution of the system state is paired with a spontaneous shift in charge distribution, and so this defined system state instantly becomes the past as a new probability density emerges. This model of Hamiltonian mechanics produces testable predictions regarding the wavelength of free energy released upon information compression. Overall, this model demonstrates how cortical neurons might achieve non-deterministic signaling outcomes through noisy coincidence detection.

## I. INTRODUCTION

Spinal reflex circuits exhibit near-perfect reliability and efficiency in transmitting information, with signaling outcomes that are easily predicted by analyzing upstream inputs [1]. By contrast, cortical neurons have highly unpredictable signaling outcomes [2]. Unlike spinal neurons, which exhibit highly deterministic firing patterns, cortical neurons allow random electrical noise to gate signaling outcomes, resulting in probabilistic firing patterns. Cortical neurons encode information through a process of noisy coincidence detection; if upstream signals and random noise temporally converge, the neuron reaches action potential threshold and opens voltage-gated sodium channels, leading to the large inward sodium ion currents which characterize the action potential [3]. Spontaneous subthreshold fluctuations in membrane potential significantly contribute to the timing of action potentials in individual cortical neurons, demonstrating that random noise plays a significant role in prompting cortical neuron signaling outcomes [4, 5]. Since electrical noise boosts error and reduces energy efficiency within digital communication channels, any well-optimized binary computing system should be highly robust to this random noise. And yet, cortical neurons actively maintain an up-state, hovering near action potential threshold and allowing noise to drive signaling outcomes [6].

The probabilistic firing patterns of cortical neurons can be modelled using Bayesian statistics [7], by introducing random connectivity [8], by employing fanofactor analysis of inter-spike variability [9], or by modifying the Hodgkin-Huxley equations to account for electrical noise [38-40]. Cognitive processes are also known to shape cortical neuron firing patterns, with contextual cues [10], prior experience [11], and expectations [12] contributing to signaling outcomes in the cortex. However, none of these methods provide mechanistic insight into how individual cortical neurons gate signaling outcomes by allowing random electrical noise to contribute to the uncertain state of each neuron. A new approach is needed.

Classically, the neuron is viewed as a binary computing unit, always in an on-state or an off-state. Here, the neuron is modeled as a two-state quantum system, with some probability of switching from an off-state to an on-state. Each ion is also superpositioned between two states, outside or inside a given neuron, with its location determined by the dynamical voltage state of each neural membrane in the vicinity and the intrinsically uncertain state of every component electron. With uncertainty being preserved rather than lost at the macro-scale, cortical neurons may be able to undergo quantum information processing. In the present model, a cortical neuron integrates probabilistic component states over some time evolution, generating a Hamiltonian operator. Since each electron in the system can contribute to the voltage state of multiple nearby neurons, the state of the entire neural network must be computed as a whole, with the system state being defined as all component pure states are transiently defined. Any redundant pure states cannot co-exist and are therefore reduced. This computational process yields eigenvalues for all state vectors and immediately restores uncertainty across the system. This theoretical model offers a mechanism by which cortical neurons might achieve non-deterministic signaling outcomes through quantum information processing.

## II. METHODS

### A. Modeling the cortical neuron as a two-state quantum system

During up-state, cortical neurons linger at their action potential threshold, allowing both upstream signals and random electrical noise to prompt a signaling outcome. So, while a neuron is classically interpreted as a binary logic gate in an ‘on’ or ‘off’ state, coded as 1 or 0, it could also be described as having some *probability* of converting to an ‘on’ state or remaining in an ‘off’ state. In this approach, a cortical neuron integrates upstream signals with random electrical noise, defining its voltage state as a function of time, as the system is perturbed. The neuron starts in off-state *ϕ*, not firing an action potential, and over time *t*, it reaches another state *χ*. And so, over some period of time, from *t*_0_ to *t*, the state of the neuron evolves from *ϕ* to *χ*. The timepath taken from one state to another is given by:

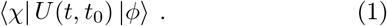

The probability of a state change can be represented in some basis:

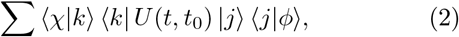

Such that *U* is completely described by base states *k* and *j*:

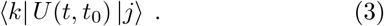

The time interval can be understood as being *t* = *t*_0_ + Δ*t*, so identifying the state of the neuron *χ* at time *t* can be understood as taking a path from one state to another:

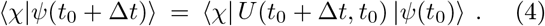

If Δ*t* = 0, there can be no state change. In this case:

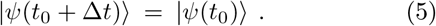

In any other case, the state of the neuron at time *t* is given by the orthonormal base states *k* and *j*, with probability amplitudes:

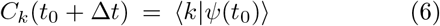

And:

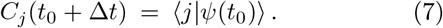

The state vector *ψ* at time *t* is a superposition of the two orthonormal base states *k* and *j*, with the sum of the squared moduli of all probability amplitudes being equal to 1:

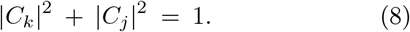

The neuronal state |*ψ*⟩ at time *t* can therefore be described as a normed state vector *ψ*, in a superposition of two orthonormal base states *k* and *j*, with probability amplitudes *C*_*k*_ and *C*_*j*_. Since the neuron starts the time evolution in state |*ψ*(*t*_0_)⟩ = *j*, its probable state at time *t* is given by:

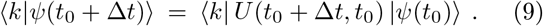

This equation can also be written in expanded form as the sum of all transition probabilities:

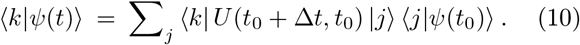

For the state vector *ψ*(*t*), the probability of a state change at time *t* is described by the U-matrix, *U*_*kj*_(*t*):

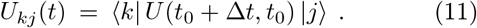

And so, all probability amplitudes are dependent on the amount of time that has passed, Δ*t*:

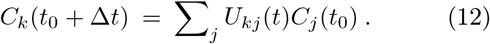

If Δ*t* = 0, there can be no state change and *k* = *j*. If Δ*t >* 0, there is some probability of a state change, where *k* ≠ *j*. As such, the two-state quantum system is described by the Kronecker delta *δ*_*kj*_:

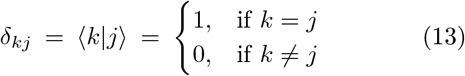

This approach describes a two-state quantum system. For small Δ*t*, each of the coefficients of the U-matrix *U*_*kj*_ differ from *δ*_*kj*_ by some amount proportional to Δ*t*, such that:

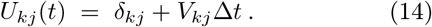

Where *V*_*kj*_ = − (*i*/*ħ*)*Ĥ*_*kj*_(*t*), with *i* representing the imaginary unit 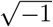 and the Hamiltonian operator term *Ĥ*_*kj*_(*t*) representing the time derivatives of each of the coefficients of *U*_*kj*_:

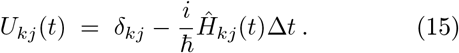

The probability amplitude *C*_*k*_(*t*) at time *t* is therefore given by:

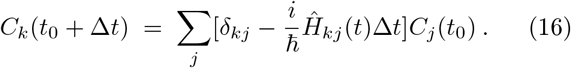

Since the sum of *δ*_*kj*_*C*_*j*_(*t*_0_) = *ψ*(*t*_0_), the latter equation simplifies to:

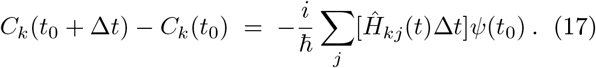

In dividing both sides of the equation by Δ*t*, it becomes apparent that any state change is a sum of all possible perturbations which affect the system under investigation, from its starting condition at time *t*_0_. Again, *C*_*k*_(*t*) is the probability amplitude ⟨*i*| *ψ*(*t*_0_) ⟩ of finding the state vector *ψ* in one of the base states *k* = *j* or *k* ≠ *j* at time *t*. The time derivative of this probability function yields the path taken:

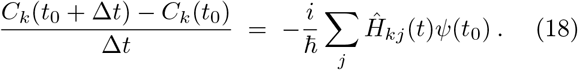

The system under consideration is therefore effectively described by the time-dependent Schrödinger equation, with any state change related to a change in energy distribution:

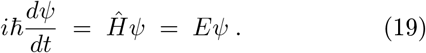

Where *i* is the imaginary unit, *ħ*= *h/*2*π* is the reduced Planck constant, *d/dt* is the amount of time that has passed since the calculation was last made, *Ĥ* is the Hamiltonian operator, which provides a vector map of all energies in the system, and *ψ* is the eigenvector describing the probable state of the quantum system after some time t has passed. This equation describes the eigenstate of a system as a function of the amount of time that has passed *t* and the amount of energy available for redistribution across the system, given by the Hamiltonian *Ĥ*. This equation allows a system state to be defined as the probability amplitude of possible paths taken since the state was last defined.

In this model, the neuronal state |*ψ*_*n*_⟩ evolves over time, with the signaling outcome at time *t* a function of the sum of all local ion states. The state of each ion in the system |*ψ*_*i*_⟩ also evolves with time, with a probabilistic location at time *t* defined in relation to every neuron, either inside that neuron or outside it. It should be noted that any change in the location of an ion is a function of the electrical fields generated by electrochemical potentials of all nearby neurons, as well as the position, momentum, and energy level of every component electron held by that ion. If uncertainty in the state of each electron is sustained for long enough to affect the behavior of the entire ion, in the presence of a dynamic electrical field, the state of each ion remains uncertain and the voltage state of each neuron therefore remains uncertain. This uncertainty is expected to be sustained in cortical neurons during an up-state, as stochastic charge flux contributes to the probability of a signaling outcome.

### B. Selecting the appropriate level of description

Each electron in the system has some possible energy state or atomic orbital *η* which is uncertain in the present moment. Any perturbances to the system, occurring over time *t*, will contribute to possible changes in the position *r* and the momentum *s* of each electron. Any energy above the electron’s ground state will contribute to the Hamiltonian and generate a complex-valued probability density describing the set of possible system states. Here, the spatial position and atomic orbital of the electron, after some amount of time has passed, are modeled as probability amplitudes distributed across five orthogonal axes, with the actual values being multiples of each Planck unit. In modeling the electron, the state vector ψ can refer to the energy state or atomic orbital, which can be any one of several orthonormal pure states |*ψ*_*x*_⟩, including a ground state and any number of unlikely excited states. The state vector *ψ* can also refer to the position *r* of the electron, across the *x, y*, and *z* axes, or the amount of time *t* that has passed to reach that new state. For example, the time-dependent Schrödinger equation for the position of a particle is:

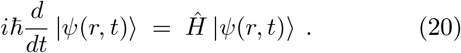

Here, the complex-valued probability amplitude can be termed as all possible positions over position space, where *R* is the position eigenvector:

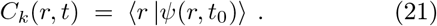

And the time-dependent Schrödinger equation for the momentum of a particle *S* is:

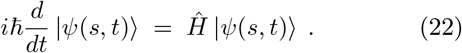

This complex-valued probability amplitude can be termed as all possible momenta over momentum space, where *S* is the momentum eigenvector:

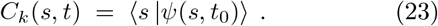

Because the state of each electron is fundamentally uncertain, the state of each ion is uncertain – particularly given the presence of a dynamic electrical field exerted by every neuron in the vicinity, as ion channels open and close and membrane potentials fluctuate. Because of this uncertainty in ion location, each ion can be considered a two-state quantum system, in relation to each neuron. A sodium ion starts the time evolution outside a given neuron, in the state *ψ*_*out*_, and has some probability of entering the neuron over time *t*. The physical location of the ion at time *t* is therefore a state vector *ψ*, in a superposition of two orthonormal base states *ψ*_*in*_ and *ψ*_*out*_, with probability amplitudes *C*_*k*_ and *C*_*j*_ such that:

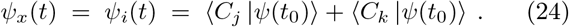

Because the state of each ion is uncertain, the state of each neuron is uncertain. Since the neuron’s membrane potential is dependent on the position of each electron in the vicinity, the neuronal voltage state is also uncertain – particularly given the fundamental uncertainty in position, momentum, and energy state of every point charge in the vicinity. Each neuron can thus be considered a two-state quantum system, with some probability of undergoing a state change over time *t*. Considering the neuron starts the time evolution at some resting potential, _*off*_, but has some probability of reaching the threshold for firing an action potential over time *t*, the state of any given neuron at time *t* is a state vector *ψ*, in a superposition of two orthonormal base states *ψ*_*on*_ and *ψ*_*off*_, with probability amplitudes *C*_*k*_ and *C*_*j*_ such that:

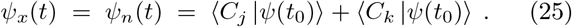

At each level of description, the quantum state can be described by the Kronecker delta. The state of the neuron at time *t* has either changed or it has not, and the position of the sodium ion at time *t* has either changed or it has not, with the possible outcomes for each computational unit in the system given by the orthonormal base states *k* and *j*, given by the Kronecker delta:

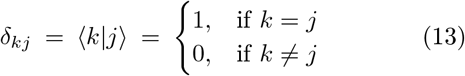

In short, the probability of a state change in a neuron, a sodium ion, or any component electron can be calculated by modeling the state vector *ψ*_*x*_. It is possible to model the entire thermodynamic system by summing the orthonormal base states of all component electrons, all component ions, or all component neurons. Each level of description can be considered a quantum system, with uncertainty preserved rather than lost at the macro-scale. To model the system at either level of description, *ρ*_*x*_ is defined as the probability of an object transitioning from one to another pure state:

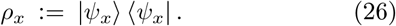

The density matrix *ρ* is comprised of an ensemble of mutually orthogonal pure states *ρ*_*x*_, each having some probability of occurring *p*_*x*_:

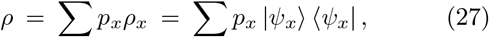

The quantity of possible states for the system is calculated by tracing the volume of probability amplitudes across a high-dimensional density matrix. This quantity is the von Neumann entropy of the system:

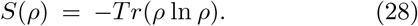

The system macrostate is described here as the mixed sum of all component pure microstates, or the mixed sum of all outer products multiplied by their transition probabilities. A state vector might represent the position, momentum, or energy state of an electron, which evolves as a function of time; the state of a given sodium ion relative to a given neuron at time *t*; or the on-off state of a given neuron at time *t*. Here, the inner product ⟨*a*_*y*_|*ψ*⟩ provides the probability that the state vector |*ψ*_*x*_⟩ assigns to the eigenvector ⟨*a*_*y*_|. As such, the probability of measuring a certain eigenvalue *a*_*y*_ equals:

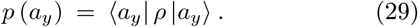

The object is in some state at time *t*_0_, and it has some probability of being found in another state at time *t*, after being transiently defined as a density of possible states. The mechanics of distributing probabilistic states into Hilbert spaces, or expending free energy to populate the Hamiltonian, allows a particle or system of particles to physically process information, then change state in a probabilistic manner. The probability of finding a system in any one eigenstate can be calculated by applying the Born rule, which equates the inner product of the state vector and its expectation value to the probability of transition to a particular actual state [13]. This rule states that any measurement of the observable has a probability *p*(*ψ*_*a*_) of being equal to an exact value *a*, at time *t* for the state vector *ψ*:

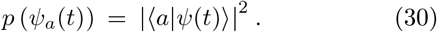

The square of the absolute value of this wavefunction is a real number, but the wavefunction itself is a complex-valued probability amplitude, which exists along an axis orthogonal to all real eigenvalues. Every probabilistic value *a* defined along this axis is dependent on the amount of time that has passed, and each of these values has a probability amplitude influenced by the constraints of the system. The primary constraint on any given electron in this system is the electrochemical potential exerted by each neuronal membrane; these membrane potentials are in turn affected by the relative position and energy state of each electron in the vicinity. To model the probabilistic behavior of the system, we will focus at the level of description of electrons, with the time-dependent state given by the Hamiltonian:

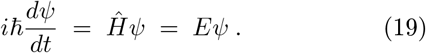

The total energy of the electron, *E*, is given by the sum of all kinetic and potential energies:

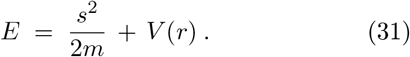

The momentum, *s*, is given by the derivative of the wavefunction with respect to position:

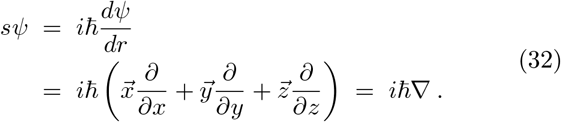

And so the square of the momentum, *s*^2^, is given by the second derivative with respect to position:

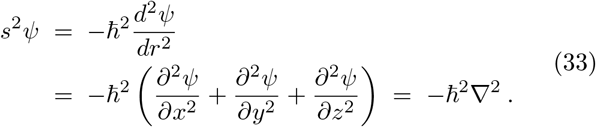

These equations can now be combined; the time-dependent state of an electron across time, across energy states, and across three spatial axes is given by a wavefunction equation that relates any time-dependent changes in the electron’s position to its momentum and energy state:

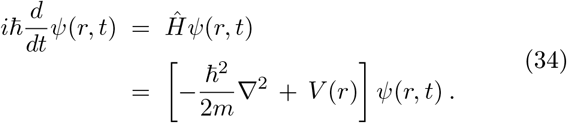

Once again, the electrochemical potential of each neural membrane is dependent upon the exact position of all electrons, in relation to the neural membrane surface, so the voltage can be modeled as the mixed sum of all component pure states. Meanwhile the exact position of each electron is dependent on the strength of these local electrical fields, with each dynamic membrane potential exerting forces that affect the position and energy state of each electron in the vicinity. This sustained uncertainty has utility in processing information, as consistencies or correlations within a temporally-bound dataset can be identified at infinitesimal timescales.

During cortical up-states, neurons actively maintain a resting potential right near the voltage threshold for firing an action potential, permitting stochastic ion leak and spontaneous membrane potential fluctuations to gate a state change in the computational unit. In the present moment, the neuron either has reached the voltage threshold to trigger a state change, or it has not; there is some probability of either outcome. The neuron remains suspended in a state of uncertainty physically encoding information, or the mixed sum of all component pure states. By sustaining uncertainty, a cortical neuron generates quantum information. These complex-valued probability amplitudes then interfere with each other, with any consistencies or correlations reducing the overall probability distribution, or the entropy of the system. This proposed biophysical process should naturally achieve a non-deterministic computation of the optimal system state in the present context.

## III. RESULTS

### A. Generating a probability density from quantum uncertainty

To understand the mechanics underlying this process, a calculation of possible system states across the neural network can be conceptualized by modeling the Poisson distribution of charge density for each electron in the system, in relation to each region of neural membrane, over some time evolution, thereby generating a sum of possible paths. Because these events do not occur with equal probability or independently of the previous system state, it is more appropriate to model the probability distribution of position and momentum vectors as *r*(*x*) and *s*(*x*), respectively, each defined across three spatial axes, with a weight function *w*(*x*) *>* 0:

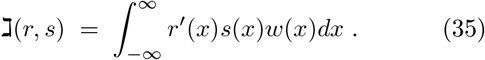

This equation yields the inner product of a Sturm-Liouville equation, with the weight function *w*(*x*) mathematically representing a quantum harmonic oscillator. This weight function embodies the quantum uncertainty of the electron state and naturally generates a Hilbert space. If this operation is symmetric or Hermitian, such that:

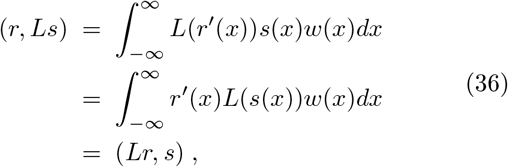

Then all polynomials will form complete sets in Hilbert space, with real eigenvalues and orthonormal eigenfunctions, providing solutions to the second-order linear differentiation equation given by:

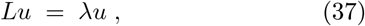

In which *λ* is a constant and *L* is a Hermitian operator defined by the real functions of *x, α, β*, and *γ*:

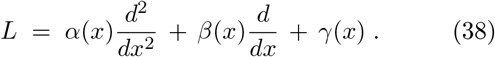

The probability of any one outcome for *r*(*x*) and *s*(*x*), thus transiently defining the system state, is given by *ρ*, the mixed sum of mutually orthogonal states *ρ*_*x*_, each occurring with some probability *p*_*x*_:

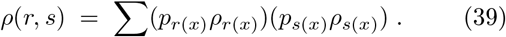

The evolution of the system state *ρ* over time *t* is given by the Liouville-von Neumann equation:

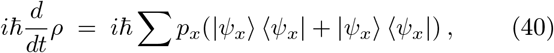

With eigenstates provided by the Hamiltonian operator and its Hermitian conjugate:

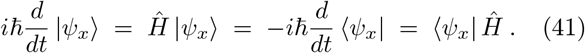

Substituting the operator and the conjugate into the Liouville-von Neumann equation yields:

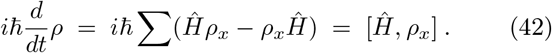

Now we can describe how the system state changes over time. A non-dissipative thermodynamic system integrates all possible microstates to generate a probability density, or a distribution of possible system macrostates. But this system does not stand alone; it interacts with the environment. All changes to the system are driven by perturbation of System A (the network of neurons or computational units) by System B (the surrounding environment), with the two systems interacting:

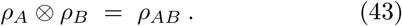

Here, System A is the cortical neural network: a non-dissipative and far-from-equilibrium system, which is perturbed by incoming caloric energy from the bloodstream and by electrochemical signals arising from the peripheral nervous system. These energetic inputs are trapped by the central nervous system, which acts as a net heat sink. Without significant energy dissipation, the particle system can use this energetic input to drive computational work, over time *t*:

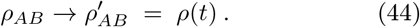

As System A is perturbed by System B, System A traps heat energy and uses it to evolve into a new state. The combined system becomes correlated, by interacting over some period of time from *ρ*(*t*_0_) to *ρ*(*t*):

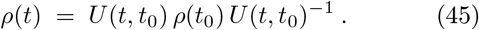

The interaction is provided by the time shift operator

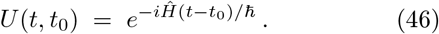

Because time and energy are intrinsically uncertain, *U* is represented by a density matrix of finite dimensionality that evolves over time, a time-dependent wavefunction, or a Hamiltonian operator which permits energy to be redistributed over time.

### B. Reducing the probability density into a single observable system state

Because the present position of a particle depends on whether it has interacted with other particles, the interdependency of eigenvectors must be taken into account here. By calculating the positions of all particles within a temperature-defined system in relation to each other, this approach ensures that particles do not turn out to occupy the same position, spin and energy state, which would render them identical. This outcome would cause the universe to lose mass, and is therefore forbidden by the Pauli exclusion principle [14]. This conservation principle can be applied to any thermodynamic system, so the total amount of energy available to the system is no more than the amount of energy stored in the system plus the net amount of energy that has entered the system over time *t*. In the central nervous system, which traps thermal energy to drive computational work, the amount of information entropy is related to the amount of free energy distributed to the Hamiltonian. Therefore, the time-dependent Schrödinger equation for an entire system of particles, which are at least partially dependent on each other, is given by the relationship between the Hamiltonian operator (the sum of all energies in the system) and the system-wide density matrix (or quantity of uncertainty) as a function of time:

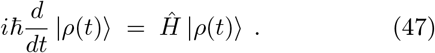

Since the probable state of each electron depends (in part) on the state of all other electrons, which are also probabilistic, the whole system must be considered together, as a sum of all pure microstates. This combined wavefunction relates the state vector *ψ* of the quantum system in relation to the passage of time *t* and the Hamiltonian operator *Ĥ*, which again corresponds to the sum of all potential energies and all kinetic energies for all particles in the system. The spectrum of the Hamiltonian operator is the set of possible outcomes at the point when the total energy of the system is measured, after some time *t* has passed. The Hamiltonian operator *Ĥ* is related to the Lagrangian function of position *r*, its time derivative 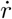 and time *t*. It is calculated by taking the Legendre transform, in order to minimize the action necessary to effect change:

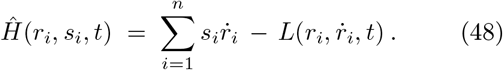

The vector spaces represented by 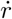and 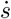 are generated by the sum of all perturbances and quantum oscillations, which are mathematically represented by the weight function within the Sturm-Liouville equation. As a result, the derivative of the Hamiltonian operator is related to any changes in position and momentum and time, and therefore can be calculated by taking the partial derivatives of these eigenvectors:

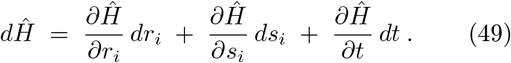

The phase space distribution *ρ*(*r, s*) describes the probability of a particular system state being selected from the total phase space volume *d*^*n*^*r d*^*n*^*s*. To describe this phase space volume, the Liouville equation yields the evolution of *ρ*(*r, s*) over time *t*:

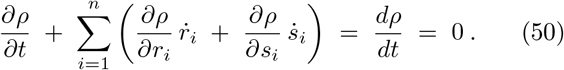

As a result of this geometrical constraint, the state vectors 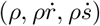 are conserved across the system, and the vector field 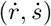 has zero divergence. This permits temperature and energy to be conserved across the system as well. The continuity equation is given by:

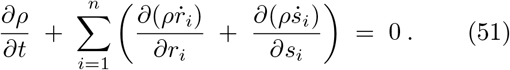

Because ∂*ρ* / ∂*t* equals zero, the continuity in the probability density can be termed as:

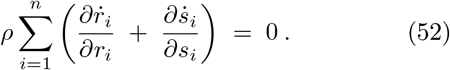

And because the vector spaces 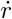 and 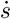 are defined as:

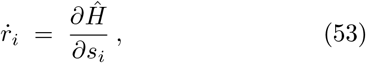

And:

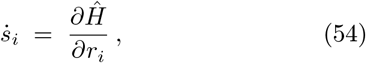

The continuity laws persist while taking the second derivative of the Hamiltonian operator. This process identifies the observable boundary of the total volume of possible system states:

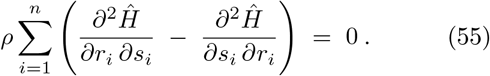

The relationship between the probability density *ρ* describing the overall system state and the Hamiltonian operator describing the total energy of the system becomes apparent in this equation, as does the relationship between position and momentum vectors and the Hamiltonian operator. Once a trace is taken across the density matrices representing *r*(*x*) and *s*(*x*), the weight function *w*(*x*) underpinning the probability distribution can be solved in relation to the other parameters.

During a system-wide computation, eigenvalues are selected and eigenstates *ψr*(*x*) and *ψr*(*x*) are transiently resolved. It should be noted that, in this formulation of temporally-proceeding events, the operators drive the computational cycle, rather than the vector states. That is, the Hamiltonian operator may evolve over time (permitting changes in position, momentum, and energy state) while the vector states themselves remain time-cindependent. Any events occurring within a time evolution, which affect the likely position or momentum of any electron, will therefore contribute to the outcome of a computation. This includes any quantum oscillations occurring over that time evolution.

By relating any probabilistic changes in the position and momentum of each electron to the distribution of energy across the system, the Hamiltonian operator effectively portrays all observable outcomes. The unitary change of basis guiding this computational process is defined by the phase factor exp(*iĤt/ħ*). Therefore, any reversal to the direction of time causes positive energies to become negative – a result which is disallowed by the first law of thermodynamics, unless the spin is also reversed with the introduction of an electric dipole moment [15]. An electric dipole moment is a spontaneous shift in the energy state of an electron in the presence of an external electric field; this event is expected to occur here, in conjunction with an abrupt reduction in probability density, as eigenvalues are calculated. If the unitary change of basis yields a zero determinant, then total dimensionality will be reduced and compatible eigenvalues will be assigned to each state vector across the entire particle system. Equivalently, taking the derivative of the entire volume of complex-valued probability amplitudes reduces all eigenstates into non-deterministic outcomes, yielding a compatible observable state on the boundary region of that high-dimensional vector space. With either description, the distribution of probabilities is reduced to a single outcome, transiently defining the system macrostate.

### C. Restoring uncertainty after the system state is transiently defined

As System A was perturbed over time, we had integrated all probabilistic component states (and the Hamiltonian operator) to create a volume of probability. So now we can take the derivative of that volume of probability (and the Hamiltonian operator) to define the surface boundary region. Doing so reduces the probability density into a single actualized state:

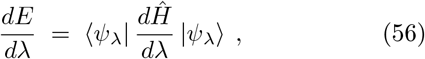

This equation is the Hellman-Feynman theorem, and it lies at the core of quantum electrodynamics [16, 17]. Here, any change in the energy state of any electron underlying the wavefunction is proportional to the change in the Hamiltonian, because the Hamiltonian corresponds to the sum total of all potential and kinetic energies in the system. It is useful to note, the total amount of energy in the system is not what is uncertain, but rather how this energy is distributed. This measure of possible system states, or energetic configurations, is the total entropy being created by the system. The total energy *E* held by the system is related to any parameter *λ* which contributes to the Hamiltonian operator, such as a shift in position, momentum, orbital number, electrical field strength, or magnetic moment. The expectation values of these observables predict how the energy in the system is distributed, to achieve an optimal system state in the present context. The derivative of the value of *E* is therefore related to the inner product of the state vectors *ψ* and the derivative of the Hamiltonian *Ĥ*, both with respect to the parameter *λ*.

Taking the derivative of the Hamiltonian means taking the derivative of all possible energy states with respect to all perturbations since the last detection event. The derivative of this eigenfunction identifies the boundary region of the high-dimensional probability density; this is equivalent to identifying the zero determinant through a unitary change of basis. In either the Hamiltonian model or the wave mechanics model or the matrix mechanics model, the complex-valued probability density is reduced and the state of each component electron in the system is transiently defined.

In developing this mathematical solution to the Schrödinger equation, Richard Feynman discovered that any wavefunction resolution has a discrete effect on the underlying particles. Once two particle systems interact, physical constraints are introduced into the combined system, and a mutually-compatible state must be found; the position and momentum and energy state of every particle become transiently defined, and all interactions within the system can therefore be deciphered using the equations of classical electrostatics, while accounting for the charge distribution across each atom in the system. But critically, the wavefunction resolution is accompanied by a discrete shift in the charge distribution across each atom in the system, in relation to their newly-defined distance from each other, locally triggering van der Waals forces [16, 17].

As a result, any perturbation to the system – for example, the movement of electrons or a shift in the local electrical field – changes the energy state of the system, in a manner related to the original unperturbed states and the derivative of the Hamiltonian. The assignment of nuclear locations, relative to other atomic nuclei in the system, prompts an alteration in the organization of electrons around the nucleus, with the charge distribution distorted from central symmetry. For two atoms interacting at a separation *D* – a distance which is large compared to the radii of the atoms – the induced dipole moment for each atom is 1*/D*^7^. The smaller the distance between atoms, the larger the dipole moment, and the larger the boost to angular momentum.

In other words, any ‘detection’ or ‘collision’ between two particle systems will cause a discrete change in component atomic states, with the more probabilistic system taking on a mutually compatible state with the more inflexible system surrounding it – allowing a probabilistic system to encode the state of its surrounding system into its own physical state. This is equivalent to a redistribution of the Hamiltonian.

Once component microstates are assigned, atoms exert effects on each other, prompting van der Waals forces between neighboring atoms [18, 19]. This is expected to be the case in the central nervous system (System A) as it encodes the state of its surrounding environment (System B). In accordance with the laws of electrodynamics, the shift in charge distribution upon wavefunction collapse should prompt van der Waals forces to emerge between sodium ions and the lipid molecules of the neural membrane. These forces may alter the likelihood of sodium ions to be transported across a nearby neural membrane. In neurons that are approaching the threshold for firing an action potential, these small currents could affect whether the neuron reaches threshold or not.

The likely position, momentum, and energy state of each electron in the system depends on the local electrochemical potential exerted by each neural membrane; this electrochemical potential in turn depends on the probable position, momentum, and energy state of each electron. The neuronal membrane potential is uncertain in the present moment, and is modeled here as the mixed sum of all component pure states.

The uncertain states populate a single density matrix or a single Hamiltonian operator, which describes all possible states of the system. The likely state of any one component electron contributes to the resolution of other component electrons, as one state constrains all other possible states [14]. Upon interaction with another particle system over some time evolution, a mutually-compatible state is identified, and the encoding system state is resolved into a singular observable outcome. The collapse of alternative eigenstates occurs as all other possible states are reduced. At that point, the information held by the system is abruptly compressed, as other possible states are eliminated. But this defined system state is transient, because it is paired with an immediate restoration of uncertainty by the alteration of charge distribution. The dipole moment induced by wavefunction resolution prompts new atomic interactions. And for successive measurements with discrete results, which do not destroy the entanglement of the particle system, each measurement with value *a* establishes the basis for a new state, which then undergoes subsequent time evolution, in accordance with the von Neumann projection postulate [20]. And so, immediately after a wavefunction is resolved, the system again begins to evolve over time, forming a new probabilistic system state.

### D. Converting probability densities to temporally-irreversible signaling outcomes

Since the expectation value of the energy state for a given particle ⟨*a*_*i*_| *ρ* |*a*_*i*_⟩ is proportional to the expectation value for its spin ⟨*ψ*| *k* |*ψ*⟩, the energy shift due to an electric dipole moment causes a sign shift in both values, and time symmetry is broken. Essentially, as ground states lose degeneracy, the resulting dipole moments will alter the attraction between sodium ions outside the neuron and atoms comprising the lipid bilayer of the neural membrane. The resulting van der Waals forces are expected to cause a permitted violation of time-reversal symmetry, thereby effecting causation within the system, as information is compressed and eigenvalues are observed.

Yet a system will only demonstrate violations of time-reversal symmetry if quantum uncertainty contributes to thermal fluctuation-dissipation dynamics. Only if coincident upstream signaling events and random electrical noise trigger a drop in membrane resistance, within the temporal parameters of ion dissipation and ion pump rectification kinetics, will inherently probabilistic events contribute to gating a neuronal state change.

This theoretical model proposes a mechanistic process by which probabilistic microstates contribute to non-deterministic signaling outcomes. Here, it is proposed that quantum uncertainty is sustained in the presence of a constantly changing electrical field, for long enough timescales to contribute to thermal fluctuations in neural membrane resistance. However, this is only expected to be the case in cortical neurons, which have been observed to gate a state change on the basis of both temporally-coincident upstream signals and random electrical noise.

### E. Conditions under which quantum fluctuations contribute to dissipation dynamics

A physical structure that actively generates an electrochemical resting potential will generate entropy and heat. The neural membrane provides resistance, and therefore some quantity of free energy will be dissipated into entropy as electrical interactions occur. However, in a heat-trapping system, this energy is not necessarily irreversibly lost; it can be used to do work [21]. The Callen-Welton fluctuation-dissipation theorem asserts that thermal fluctuations drive a response function affecting the impedance of the structure, given by *χ*(*t*):

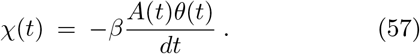

Where *β* = *k*_*B*_*T, θ*(*t*) is the Heaviside step function, and *A*(*t*) is the expectation value of *O*_*k*_(*t*), an observable that is subject to thermal fluctuations in a dynamical system. If decoherence timescales within the system are longer than the timescales of ion dissipation and ionization dynamics, quantum uncertainty can materially contribute to ion behavior at the neuronal membrane [22]. That is, if the time-dependent perturbation in the behavior of *ions* relies on any quantum fluctuations in the position, momentum, or energy state of component *electrons*, then the ensemble average of each observable (a measure of fluctuation, given by the Hermitian operator [*O*_*k*_(*t*), *O*_*j*_(*t*_0_)]) will be related to the response function (a measure of dissipation, given by *χ*(*t* − *t*_0_), as a function of time. This relationship is given by the Kubo fluctuation-dissipation formula [23, 24]:

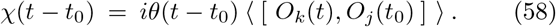

This response function, *χ*(*t*), can be written as a function of oscillatory events:

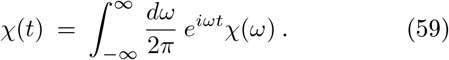

If *t* < 0, then *iωt* will be negative and *e*^*iωt*^ will be zero, so the entire response function *χ*(*t*) will be zero. As such, the quantum states underpinning this spectral function can only causally contribute to dissipation dynamics as time moves in a forward direction. The Fourier transform of the response function provides for dissipation and fluctuation dynamics in the frequency domain, given by *χ*(*ω*):

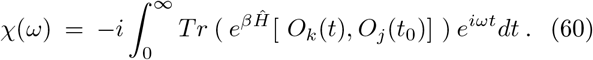

This time-dependent function provides the density of possible states for a particle system, which emerge in the presence of a perturbation or changing electrical field. And so [*O*_*k*_(*t*), *O*_*j*_(*t*_0_)] is simply a description of how the density matrix, the wavefunction, or the Hamiltonian operator changes over some time evolution, compared with the probability that all energy states would rearrange in that same way under time reversal. As such, the response function *χ*(*ω*) can be written in expanded form:

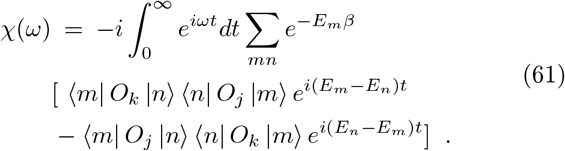

If the uncertainty in electron position, momentum, and energy state is sustained in the presence of a constantly changing electrical field, then quantum fluctuations may contribute to ion behavior, affecting the dynamics of ions interacting with the electrochemical potential of the neural membrane. In this case, quantum fluctuations may contribute to the probability of a state change in the computational unit, from an off-state to an on-state. However, the state of each neuron at the moment the Hamiltonian operator is resolved will govern its response. Neurons in a cortical up-state, which allow stochastic events to gate a signaling outcome, may be nudged toward action potential threshold or away from it as uncertainty is abruptly reduced. Indeed, *only* neurons in cortical upstate, allowing random noise to gate a signaling outcome, will be nudged to fire a signal. Meanwhile, neurons receiving suprathreshold stimulation will exhibit deterministic firing patterns, and neurons receiving insufficient upstream inputs will not be triggered to fire at all.

### F. The expected wavelength of spontaneous free energy release

If cortical neural networks engage in ambient-temperature quantum computation, then these far-from-equilibrium thermodynamic systems must bidirectionally exchange free energy for information, with any free energy expended on information generation during the initial stage of the thermodynamic computing cycle being partially recovered during the information compression stage. As such, discrete quantities of free energy should be released, local to any reduction of uncertainty, as an optimal system state is selected from some probability distribution. Any thermal fluctuation should correspond to a shift in electron energy states, boosting the angular momentum of individual ions. These events should therefore be observable. To evaluate this hypothesis, we can calculate the expected effects of quantum fluctuation-dissipation dynamics in the mammalian central nervous system.

If the perturbation to the state of an ion during some time evolution relies on any random changes to the spin or energy state of a component electron, then the ensemble average of each observable (a measure of fluctuation, given by 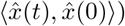) will be related to the response function (a measure of dissipation, given by *χ*(*t*)) in the frequency domain [23]. This relationship essentially models how the oscillatory behavior of a quantum system, along an imaginary axis, affects the thermal dynamics of the observable system:

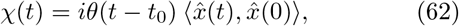

Where *θ*(*t*) is the Heaviside step function and the response *χ*(*t*) provides an expectation value for *x*(*t*), which is a time-dependent ‘observable’ subject to thermal fluctuation in a dynamical system. The time-dependent equation is given by:

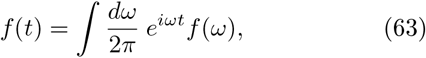

And its Fourier transform is given by:

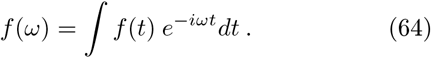

If the eigenvalues are to be real, the sum of the real part and the imaginary part of the response function over some time evolution *χ*(*t* − *t*_0_) must also be real. The full response function is given by:

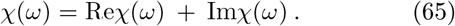

The real part of the response function is given by:

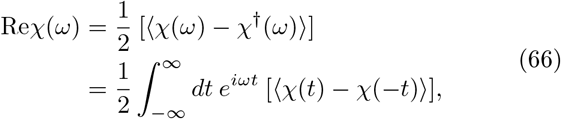

And the imaginary part of the response function is given by:

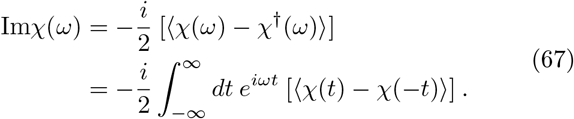

The energy of quantum fluctuations *E* (*ω*) is related to the frequency *ω*:

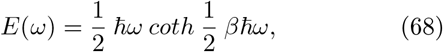

Where *β* = 1*/k*_*B*_*T*. The classical power spectrum is related to the complex-valued quantum spectral density in such a way that quantum noise can contribute to the local thermal density under certain conditions:

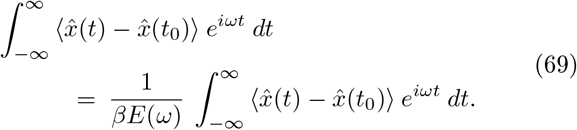

If *E* > *k*_*B*_*T*, with a high temperature, a broad distribution of electrons across energy states, and low occupancy of energy states, then the quantum contribution to ion behavior is negligible, and the behavior of ions will reduce to Boltzmann-Maxwell statistics. If *E* < *k*_*B*_*T*, with electrons at low frequencies obeying the Rayleigh-Jeans law, then the quantum contribution to ion behavior is also negligible, and again the behavior of ions will reduce to classical Boltzmann-Maxwell statistics. Only in cases of high particle density, when the energy held by an ion is greater than its chemical potential, and *E* > *k*_*B*_*T*, will quantum fluctuations contribute to ion behavior. This is predicted to be the case in the mammalian brain.

If quantum fluctuations do indeed contribute to ion behavior in biological systems, then *E* must be greater than *k*_*B*_*T*. Since:

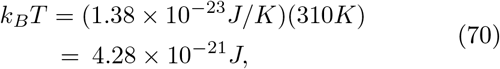

And:

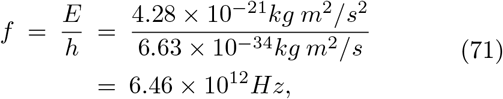

it is expected that high-energy particles of *E* > *k*_*B*_*T* will be observed in the central nervous system at 37° C (310K). Specifically, these high-energy particles should have a frequency of *f* > 6.46 × 10^12^ Hz, a wavelength of *λ* < 46 microns, or an energy of *E* > 0.0267 eV, within the infrared light spectrum. Spontaneous emissions of photons in this range have indeed been observed in mammalian brain tissue [25-28] and infrared stimulation of the brain has been shown to have a functional effect on neural activity [29-31]. Further studies are needed to measure the exact wavelengths of these photon emissions and temporally correlate these events with neuronal signaling outcomes.

In summary, it is predicted that photons, specifically in the infrared range of the electromagnetic spectrum, should be released upon information compression in the mammalian brain. Since the first law of thermodynamics states that energy cannot be created nor destroyed, any system capable of reducing entropy to achieve a non-deterministic computation must release free energy upon information compression. If quantum computing does occur in cortical neural networks, then spontaneous thermal fluctuations should be observed, locally to any reduction of uncertainty. These thermal fluctuations are expected to drive synchronous firing across the neural network.

Therefore, in this approach, the reduction of thermodynamic entropy is paired with *both* the selection of an optimal system state from a large probability distribution (one that is thermodynamically favored to correlate with the surrounding environment) *and* the release of thermal free energy (which is used to physically instantiate the solution to that computational problem).

### G. Specific predictions of this model

If cortical neural networks are indeed quantum computing systems rather than classical computing systems, then evidence of quantum information generation and compression should be observed in the neocortex. As such, this theory makes specific predictions for cortical neurons, with regard to coulomb scattering and decoherence timescales [32]. This theory also makes specific predictions about the expected effects of electromagnetic stimulation and various pharmacological interventions in cortical neural networks [33]. Some additional specific predictions of the theoretical framework, prompted by the present model, include:

#### 1. Thermal free energy is spontaneously released during the computation, as information is compressed

Infrared particles with wavelengths of *λ* < 46 microns or *f* > 6.46 × 10^12^ Hz should spontaneously appear at the neural membrane during cortical information processing. This prediction can be tested with sensitive infrared detection devices rather than classical electrodes or imaging systems; the spontaneous release of infrared-wavelength particles should be observed in the brain as uncertainty is resolved into signaling outcomes. A quantitative increase in these particles should be observed, for example, upon perceptual recognition of a highly uncertain visual or auditory stimulus, with a strong temporal correspondence to P300 event-related potentials in the cerebral cortex. By contrast, this spontaneous thermal free energy release should not occur in the case of an epileptic seizure - when constitutive ion channel activation, rather than information processing, leads to highly synchronized neural activity across the cerebral cortex. Of course, spontaneous emissions of photons in this range have been observed in mammalian brain tissue [25-28], and infrared stimulation of the brain has been shown to have a functional effect on neural activity [29-31], but further studies are needed to measure the exact wavelengths of these photon emissions and temporally correlate these events with neuronal signaling outcomes.

#### 2. The spontaneous release of thermal free energy during information compression prompts synchronized firing across the neural network

This theoretical framework describes both a computational process and a thermodynamic process, since information compression is both the selection of an optimal system state from a large probability distribution and the reduction of thermodynamic entropy. This system-wide computational event, resolving the uncertainty in component pure states, should lead to a spontaneous release of free energy, driving synchronous firing of neurons across the network. Synchronous neural activity is indeed observed at a range of frequencies in cortical neural networks, and is considered a correlate of higher-order cognitive processes [34, 35]. However, the high frequency oscillations observed during perceptual tasks cannot be modeled by coupling and recruitment under classical assumptions and timescales [36, 37]. Here, information compression events are predicted to prompt spontaneous synchronized activity across sparsely-distributed neuron populations. This coordinated activity is predicted to occur in cells that allow random noise to gate signaling outcomes (e.g. cortical neural circuits) but not in cells that act entirely deterministically (e.g. spinal reflex circuits). While this oscillatory activity, occurring at a range of nested frequencies, has been observed in the mammalian brain, additional studies could explore the potential correlation between probabilistic coding and network-level activity in avian and cephalopod species. If synchronous activity is caused by classical methods of signal propagation, rather than being the result of a system-wide non-deterministic computation, then both classical simulations of cortical neural networks and spinal reflex circuits should readily demonstrate fast and slow oscillations. If instead synchronous activity across the network is caused by information compression, paired with free energy release, then synchronous firing should be eliminated by absorption of the predicted wavelengths and should be prompted by introduction of these wavelengths.

#### 3. Higher brain temperatures should lead to higher-amplitude EEG signals, as well as a qualitatively richer perceptual experience

Thermoregulatory control is a key requirement for thermodynamic computation, with higher temperatures providing more energy for the Hamiltonian operator. For this reason, higher temperatures – such as those generated by fever – should lead to an increase in the distribution of possible macrostates for the neural network, correlated with an increase in the richness and diversity of information content. This uncertainty should also lead to greater difficulty in effectively compressing the information and higher amplitude EEG activity as information is compressed. Given an uncertain stimulus, it should be more difficult to reduce the system-wide probability distribution into a single outcome, causing perceptual errors. Therefore, this model predicts that a raised temperature within the central nervous system should lead to increased vividness in perception and decreased accuracy during time-constrained information processing. Under classical assumptions, high temperatures may lead to bodily dehydration or protein degradation, leading to errors in perception due to the unavailability of biochemical resources. In this model, thermal free energy availability directly contributes to information generation, so increasing this thermodynamic quantity should immediately increase the amount of information available to the system to be perceived. If there are time constraints for making a decision, in the context of a highly uncertain stimulus, there should be more frequent errors at higher temperatures, since more information must be parsed.

#### 4. Lower brain temperatures should lead to lower-amplitude EEG signals, as well as a diminished perceptual experience and impaired cognitive function

A difficulty maintaining sufficient temperature across the central nervous system leads to a decrease in effective information processing within the neural network. Under classical assumptions, low temperatures lead to a decrease in blood flow and oxygen availability, leading to a loss of consciousness due to this biological shortfall. In this model, reduced thermal free energy is predicted to directly reduce the amount of information content generated, causing percepts to become dim or degraded. The reduced information content should be correlated with decreased amplitude in event-related potentials associated with percept recognition. This model also predicts that if the thermal free energy released during information compression is dissipated too quickly, due to impaired thermoregulatory control or metabolic inefficiency, this quantity will be lost before it can be productively directed toward work. Thus, thermodynamic inefficiency should lead both to impairments in synaptic remodeling during learning and to impairments in directing appropriate motor output in response to sensory input. As such, people with reduced metabolic efficiency and an inability to maintain temperature, such as aged individuals, are predicted to experience diminished perception, decreases in memory consolidation, and a reduced ability to initiate voluntary behavior. Maintaining a stable brain temperature is therefore predicted to promote neurological health and stave off dementia.

## IV. DISCUSSION

The neuron is classically viewed as a transistor, always in either an on-state or an off-state. Here, the cortical neuron is modeled as a qubit, with some probability of transitioning from an off-state to an on-state over some time evolution. In this new approach, a state change in the computational unit relies on inherently probabilistic events; this model specifically takes into account the contribution of random electrical noise in gating cortical neuron signaling outcomes.

With this new approach, each electron has a range of possible spatial locations and energy states, in relation to each neural membrane, after some amount of time has passed. These complex-valued probability amplitudes, or component pure states, sum together to create a complex-valued probability density. This mixed sum of component microstates is the physical definition of quantum information, or von Neumann entropy. Some neurons may receive sufficient upstream signals to push them over action potential threshold; others will be more uncertain, and sensitive to the contribution of random noise. Using the well-established toolkits and mathematical formalism of Hamiltonian mechanics, this study demonstrates how cortical neurons retaining a state of uncertainty could physically generate and compress information to achieve non-deterministic signaling outcomes.

In accordance with the laws of mechanics, inherently probabilistic component states are integrated to populate a Hamiltonian operator. The Hamiltonian operator is then differentiated with respect to all perturbations to the system. The redistribution of energy that results from this computational process assigns eigenvalues for the spatial location and atomic orbital of each electron in the system at a single point in time. This newly-actualized system state immediately becomes the past, as new probability amplitudes emerge to describe the likely position, momentum, and energy state of each electron in the present moment, in relation to the electrochemical potential exerted by each nearby neural membrane surface. There are two additional ways to describe this computational process, provided in sister reports:

In accordance with the laws of thermodynamics, free energy must be expended to create information and this free energy is partially recovered upon information compression. In this model, probabilistic component pure states can be represented algebraically by a density matrix [32]. The density matrix undergoes a unitary change of basis, as the system state is perturbed by its surrounding environment over some time evolution. The diagonalization of the density matrix yields a zero determinant, leading to observables on the boundary region of that high-dimensional probability density.

In accordance with the laws of holography, these probabilistic component pure states can also be represented geometrically as complex-valued waves or wavefunctions [33]. These complex-valued wavefunctions or probability amplitudes constructively and destructively interfere on the charge-detecting polymer surface of the neural membrane. As a result of this physical interference between probability amplitudes, dominant probabilities are favored, becoming actualized on the boundary region of the high-dimensional probability density.

This model of Hamiltonian mechanics is complemented by these models of matrix mechanics and wave mechanics, which also demonstrate a computational process of information generation and compression. In short, cortical neurons may be better described as qubits, encoding von Neumann entropy, rather than classical bits, encoding Shannon entropy. This theoretical framework for non-deterministic computation at ambient temperatures may not only provide useful insight into the operation of biological systems, but also drive advances in machine learning and decision-making.

## ACKNOWLEDGMENTS

The author received support for this work from the Western Institute for Advanced Study, with generous donations from Jason Palmer, Bala Parthasarathy, and Vanguard Charitable.

## References

[1] Powers RK, Binder MD. (1995) Effective synaptic current and motoneuron firing rate modulation. J Neurophysiol 74(2): 793–801.

[2] Softky, W.R. and Koch, C.(1993) The highly irregular firing of cortical cells is inconsistent with temporal integration of random EPSPs. J Neurosci 13(1): 334–50.

[3] Armstrong, C.M. and Hille, B. (1998) Voltage-gated ion channels and electrical excitability. Neuron 20(3): 371–80.

[4] Stern, E.A., Kincaid, A.E., and Wilson, C.J. (1997) Spontaneous subthreshold membrane potential fluctuations and action potential variability of rat corticostriatal and striatal neurons in vivo. J Neurophysiol 77(4): 1697–715.

[5] Dorval, A.D. and White, J.A. (2005) Channel noise is essential for perithreshold oscillations in entorhinal stellate neurons. J Neurosci 25(43): 10025–8.

[6] Haider, B., et al. (2006) Neocortical network activity in vivo is generated through a dynamic balance of excitation and inhibition. J Neurosci 26(17): 4535–45.

[7] Beck, J.M., et al. (2008) Probabilistic population codes for Bayesian decision making. Neuron 60(6): 1142–52.

[8] Maoz, O., et al. (2020) Learning probabilistic neural representations with randomly connected circuits. Proc Natl Acad Sci U S A 117(40): 25066–73.

[9] Fayaz, S., Fakharian, M.A., and Ghazizadeh, A. (2022) Stimulus presentation can enhance spiking irregularity across subcortical and cortical regions. PLOS Comput Biol 18(7):e1010256.

[10] Zhou, H., et al. (2020) Spatiotemporal dynamics of brightness coding in human visual cortex revealed by the temporal context effect. Neuroimage 205: 116277.

[11] Snyder, J.S., et al. (2015) How previous experience shapes perception in different sensory modalities. Front Hum Neurosci 9: 594.

[12] Kok, P., van Lieshout, L.L., and de Lange, F.P. (2016) Local expectation violations result in global activity gain in primary visual cortex. Sci Rep 6:37706.

[13] Born, M. (1955) Statistical Interpretation of Quantum Mechanics. Science 122(3172): 675–79.

[14] Pauli, W. (1925) Über den Zusammenhang des Abschlusses der Elektronengruppen im Atom mit der Komplexstruktur der Spektren. Zeitschrift für Physik 31: 765–83.

[15] Khriplovich, I.B. and Lamoreaux, S.K. (2012) CP Violation Without Strangeness: Electric Dipole Moments of Particles, Atoms, and Molecules. Berlin: Springer

[16] Feynman, R.P. (1939) Forces in Molecules. Phys Rev 56: 340.

[17] Esteve, J.G., Falceto, F., and Garcia Canal, C. (2010) Generalization of the Hellmann-Feynman theorem. Phys Lett A 374(6): 819–22.

[18] Bethe, H.A. (1947) The electromagnetic shift of energy levels. Phys Rev 72(4): 339.

[19] Holstein, B.A. (2001) The van der waals interaction. Am J Phys 69(4): 441–9.

[20] von Neumann, J. (1932) Mathematical Foundations of Quantum Mechanics. Berlin: Springer 205: 116277.

[21] Callen, H.B. and Welton, T.A. (1951) Irreversibility and generalized noise. Phys Rev 83: 34.

[22] Tegmark, M. (2000) Why the brain is probably not a quantum computer. Information Sciences 128: 155–79.

[23] Kubo, R. (1966) The fluctuation-dissipation theorem. Rep Prog Phys 29: 255–84.

[24] Martyushev, L.M., Nazarova, A.S. and Seleznev, V. D. (1966) On the problem of minimum entropy production in the nonequilibrium stationary state. J Phys A 40(3):371–80.

[25] Isojima, Y., Isoshima, T., Nagai, K., Kikuchi, K., and Nakagawa, H. (1995) Ultraweak biochemilumines-cence detected from rat hippocampal slices. Neuroreport 6(4):658–60.

[26] Kobayashi, M., Takeda, M., Sato, T., Yamazaki, Y., Kaneko, K., Ito, K., Kato, H., and Inaba, H. (1999) In vivo imaging of spontaneous ultraweak photon emission from a rat’s brain correlated with cerebral energy metabolism. Neurosci Res 34(2):103–13.

[27] Kataoka, Y., Cui, Y., Yamagata, A., Niigaki, M., Hirohata, T., Oishi, N., and Watanabe, Y. (2001) Activity-dependent neural tissue oxidation emits intrinsic ultraweak photons. Biochem Biophys Res Commun 285(4):1007–11.

[28] Tang, R. and Dai, J. (2014) Spatiotemporal imaging of glutamate-induced biophotonic activities and transmission in neural circuits. PLoS One 9(1):e85643.

[29] Amaroli, A., et al. (2018) Near-infrared laser photons induce glutamate release from cerebrocortical nerve terminals. J Biophotonics 11(11):e201800102.

[30] Naeser, M.A., Ho, M.D., Martin, P.I., Hamblin, M.R., and Koo, B.B. (2020) Increased functional connectivity within intrinsic neural networks in chronic stroke following treatment with red/near-infrared transcranial photobiomodulation. Photobiomodul Photomed Laser Surg 38(2):115–131.

[31] Tan, X., Rajguru, S., Young, H., Xia, N., Stock, S.R., Xiao, X., Richter, C.P. (2015) Radiant energy required for infrared neural stimulation. Sci Rep 5:13273.

[32] Stoll, E.A. (under review) Random electrical noise drives non-deterministic computation in cortical neural net-works.

[33] Stoll, E.A. (under review) Modeling electron interference at the neuronal membrane yields a holographic projection of representative information content.

[34] Buzsaki, G. and Draguhn, A. (2004) Neuronal oscillations in cortical networks. Science 304(5679):1926–1929.

[35] Engel, A.K. and Singer, W. (2001) Temporal binding and the neural correlates of sensory awareness. Trends Cogn Sci 5(1):16–25.

[36] Stacey, W.C., Krieger, A., and Litt, B. (2011) Network recruitment to coherent oscillations in a hippocampal computer model. J Neurophysiol 105(4):1464–81.

[37] Whittington, M.A., Cunningham, M.O., LeBeau, F.E.N., Racca, C., and Traub, R.D. (2010) Multiple origins of the cortical gamma rhythm. Dev Neurobiol 71(1):92–106.

[38] Rinzel, J. and Miller, R.N. (1980) Numerical calculation of stable and unstable periodic solutions to the Hodgkin-Huxley equations. Math Biosci 49:27–59.

[39] Rowat, P. (2007) Interspike interval statistics in the stochastic Hodgkin-Huxley model: coexistence of gamma frequency bursts and highly irregular firing. Neural Comput 19(5):1215–1250.

[40] Austin, T.D. (2008) The emergence of the deterministic Hodgkin-Huxley equations as a limit from the underlying stochastic ion channel mechanism. Ann Appl Prob 18:1279–1325.

